# Affinity sedimentation and magnetic separation with plant-made immunosorbent nanoparticles for therapeutic protein purification

**DOI:** 10.1101/2021.11.05.467285

**Authors:** Matthew J. McNulty, Anton Schwartz, Jesse Delzio, Kalimuthu Karuppanan, Aaron Jacobson, Olivia Hart, Abhaya Dandekar, Anatoli Giritch, Somen Nandi, Yuri Gleba, Karen A. McDonald

## Abstract

The virus-based immunosorbent nanoparticle is a nascent technology being developed to serve as a simple and efficacious agent in biosensing and therapeutic antibody purification. There has been particular emphasis on the use of plant virions as immunosorbent nanoparticle chassis for their diverse morphologies and accessible, high yield manufacturing via crop cultivation. To date, studies in this area have focused on proof-of-concept immunosorbent functionality in biosensing and purification contexts. Here we consolidate a previously reported pro-vector system into a single *Agrobacterium tumefaciens* vector to investigate and expand the utility of virus-based immunosorbent nanoparticle technology for therapeutic protein purification. We demonstrate the use of this technology for Fc-fusion protein purification, characterize key nanomaterial properties including binding capacity, stability, reusability, and particle integrity, and present an optimized processing scheme with reduced complexity and increased purity. Furthermore, we present a coupling of virus-based immunosorbent nanoparticles with magnetic particles as a strategy to overcome limitations of the immunosorbent nanoparticle sedimentation-based affinity capture methodology. We report magnetic separation results which exceed the binding capacity of current industry standards by an order of magnitude.

## 1. Introduction

Virus-based nanomaterials are proving to be uniquely accessible, precise, and efficacious solutions to problems in fields ranging from energy to medicine (Wen and Steinmetz, 2016). Plant viruses serve as a particularly interesting biologically-derived nanomaterial for their inherent advantages of host specificity-related human safety (Nikitin *et al.*, 2016), simplicity of *in planta* cultivation (Hefferon, 2017), and wide variety of particle architectures and functionalities (Ibrahim *et al.*, 2019). Plant viral nanoparticles and virus-like particles have been studied for diverse biotechnical applications including gene therapy (Azizgolshani *et al.*, 2013; Czapar and Steinmetz, 2017), vaccines (Balke and Zeltins, 2019; Canizares *et al.*, 2005), medical imaging (Aljabali *et al.*, 2019; Shukla and Steinmetz, 2015b), drug delivery (Bruckman *et al.*, 2018; Lebel *et al.*, 2016), and biosensors (Bäcker *et al.*, 2016; Soto *et al.*, 2006).

The concept of a plant virus-based immunosorbent nanoparticle (VIN), a plant virus or virus-like particle displaying antibody-binding proteins, has been proposed to capture antibodies for biosensing (Kuo *et al.*, 2018; Uhde-Holzem *et al.*, 2016) and therapeutic antibody purification (Werner *et al.*, 2006). This nascent technology is one approach to address the need to reduce capital intensity for equitable and accessible antibody-related healthcare solutions, which could be harnessed to treat more prevalent diseases with availability of inexpensive and adequate production and purification capacity (Buyel *et al.*, 2017). Purification can cost up to 80% of the total manufacturing expenses for antibody and other biopharmaceutical products (Yang *et al.*, 2020). The simple and bioregenerable VIN technology is also one that may transcend terrestrial needs as humankind considers extended duration space exploration and is faced with stringent life support system requirements in perhaps the most limited resource environment that humans will face (Aglietti, 2020; Menezes *et al.*, 2015). Recent literature highlights the potential of plant-based manufacturing to close human health risk gaps for manned exploration missions (Matthew J. McNulty *et al.*, 2021).

Initial VIN research has primarily focused on nanomaterial design, considering three plant virion chassis (potato virus X (Uhde-Holzem *et al.*, 2016), bamboo mosaic virus (Kuo *et al.*, 2018), turnip vein clearing virus (TVCV) (Werner *et al.*, 2006)) and several ligand display strategies, including multiple fusion sites on the coat protein, linker inclusions, modulations of ligand display density, and two different immunosorbent ligands (both based on functional fragments of *Staphylococcus aureus* Protein A). Additional research is needed to evaluate broader functionalities of VIN technology, characteristics for reliable biomanufacturing, and compatibility with advanced multi-material configurations.

There have been multiple approaches to engineering virus-based nanomaterials into multi-material configurations including layer-by-layer assembled thin biofilms (Tiu *et al.*, 2017), electrospun nanofibers (Shin *et al.*, 2014), and bio-functionalized magnetic particles, to name a few. Within these approaches, bio-functionalized magnetic particles have been distinguished at large as an important platform within biosensing (Zhang and Zhou, 2014), and protein purification (Schwaminger *et al.*, 2019). However, the virus-based nanomaterial research exploring bio-functionalized magnetic particles to date has been limited to gene therapy (Chan *et al.*, 2005; Majidi *et al.*, 2015) and molecular imaging (Huang *et al.*, 2011; Shukla and Steinmetz, 2015a). Given the demonstrated ability of virus-based nanomaterials as reagents to enhance target binding and sensitivity over traditional ligands (Koch *et al.*, 2015; Sapsford *et al.*, 2006; Soto *et al.*, 2009) we perceive a general synergy and advantage in developing virus-functionalized magnetic particles for sensing and protein purification.

In this study we present a new vector for production of VINs, develop an optimized purification process for VINs that is generalizable to other plant virus-based nanomaterials, characterize key functional VIN properties, and in the process, identify potential limitations of the VIN methods used to date. In response to identification of these limitations, we present a novel VIN-magnetic particle coupled system to overcome these limitations. Preliminary results suggest enhanced immunosorbent characteristics as compared to commercial immunosorbent magnetic particle standards and also provide a new perspective for utilization of plant virus-based nanomaterials.

## 2. Results

### 2.1. Production of a plant virus-based immunosorbent nanoparticle

Intact VINs consisting of an assembled tobamovirus, TVCV, presenting a C-terminal coat protein fusion to a flexible linker domain (GGGGS)_3_ coupled to a *S. aureus* Protein A fragment (domains D and E) were successfully produced in *Nicotiana benthamiana* plants via agroinfiltration and subsequently purified to a moderate extent (Figure 1a – d). An illustration of the construct schematic and results of the PCR and DNA sequence verification of the transformation are included in Supporting figures: Figure S1, Figure S2 and Supporting table: Table S1. The vector used here simplifies previously published *A. tumefaciens* vectors (Werner *et al.*, 2006) by combining multiple provectors into a single vector capable of producing intact VINs.

**Figure 1.**
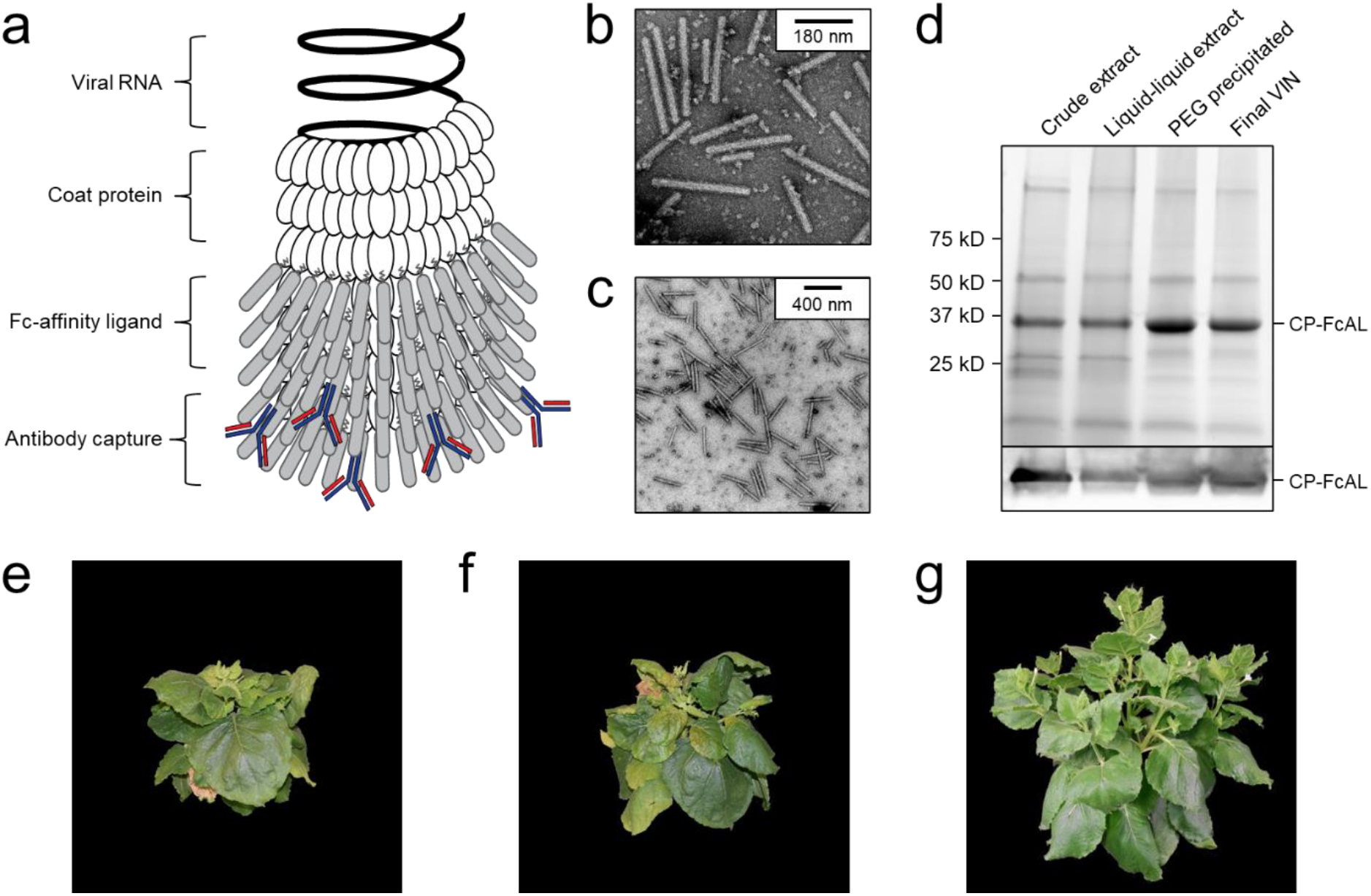
Production of a plant virus-based immunosorbent nanoparticle (VIN). (a) An illustrative depiction of a VIN. The native plant virus (viral nucleic acid encapsulated by coat protein) is fused via a peptide linker to an Fc-affinity ligand, which confers immunosorbent functionality to the plant virus. (b, c) Negative stain transmission electron microscope images of VIN in crude plant extract solution produced via agroinfiltration in *N. benthamiana* plants. (d) Reducing condition SDS-PAGE (upper) and Western blot (lower) of the VIN preparation marked at the band height corresponding to VIN coat protein Fc-affinity ligand fusion (CP-FcAL). Representative photographs of 7-week-old *N. benthamiana* plants incubated in a controlled environment facility for 14 days post-infection with mechanically transmitted (e) VIN (produced using vector pICH25892), and (f) wild-type tobacco mosaic virus, as compared to (g) uninfected healthy plants.

Agroinfiltrated *N. benthamiana* plants showed signs of viral infection typical of tobamoviruses (yellowing of leaves, stunted growth; data not shown). The coat protein fusion was expressed at high levels in *N. benthamiana* plants collected 6-14 days post-infiltration. Furthermore, transmission electron microscope (TEM) images show that fully assembled virion particles were formed (Figure 1b-c).

SDS-PAGE results confirm that there is a band at the expected size of the VIN coat protein Fc-affinity ligand fusion (CP-FcAL) (~33.5 kD) and Western blot results confirm that it is an identity match for the expected CP-FcAL (via anti-protein A antibody). We did not observe bands corresponding to unfused Fc-affinity ligand on SDS PAGE gels or in Western blots, although SDS-PAGE results do present the possibility of a minor presence of CP-FcAL degradation products. The CP-FcAL protein identity was also confirmed using mass spectrometry (Supporting figure: Figure S3).

In addition to agroinfiltration, we also demonstrated that mechanical transmission using the VINs generated by agroinfiltration is a viable route for production of fully assembled and functional plant virus-based immunosorbent particles. Mechanical transmission of the solution containing fully assembled VINs yielded systemic plant infection and morphological change (Figure 1e – g), indicating that the VINs retain systemic mobility with the immunosorbent fusion protein. Agroinfiltration-based VIN expression induced comparable *N. benthamiana* plant morphology (data not shown).

### 2.2 Fc-protein capture and elution

Protein A is well known to bind strongly with the conserved fragment crystallizable (Fc) region of many species and subclass variants of IgG. We show that VINs retain general immunosorbence for several species and subclasses of IgG (Supporting figure: Figure S4).

Next, we demonstrate that VINs are capable of capturing and then eluting human immunoglobulin G (hIgG) using a low pH elution mechanism (Figure 2a – b). VINs produced using mechanical transmission were also shown to retain immunosorbent functionality (Supporting figure: Figure S5). Tests using bovine serum albumin (BSA) as the target capture protein confirm that sedimentation of the target protein, and largely that of the VIN, required specific binding interactions (Figure 2c). It was also observed that VINs would sediment in the absence of binding target proteins at centrifugation of 20,000 x g for 90 minutes (Supporting figure: Figure S6).

**Figure 2.**
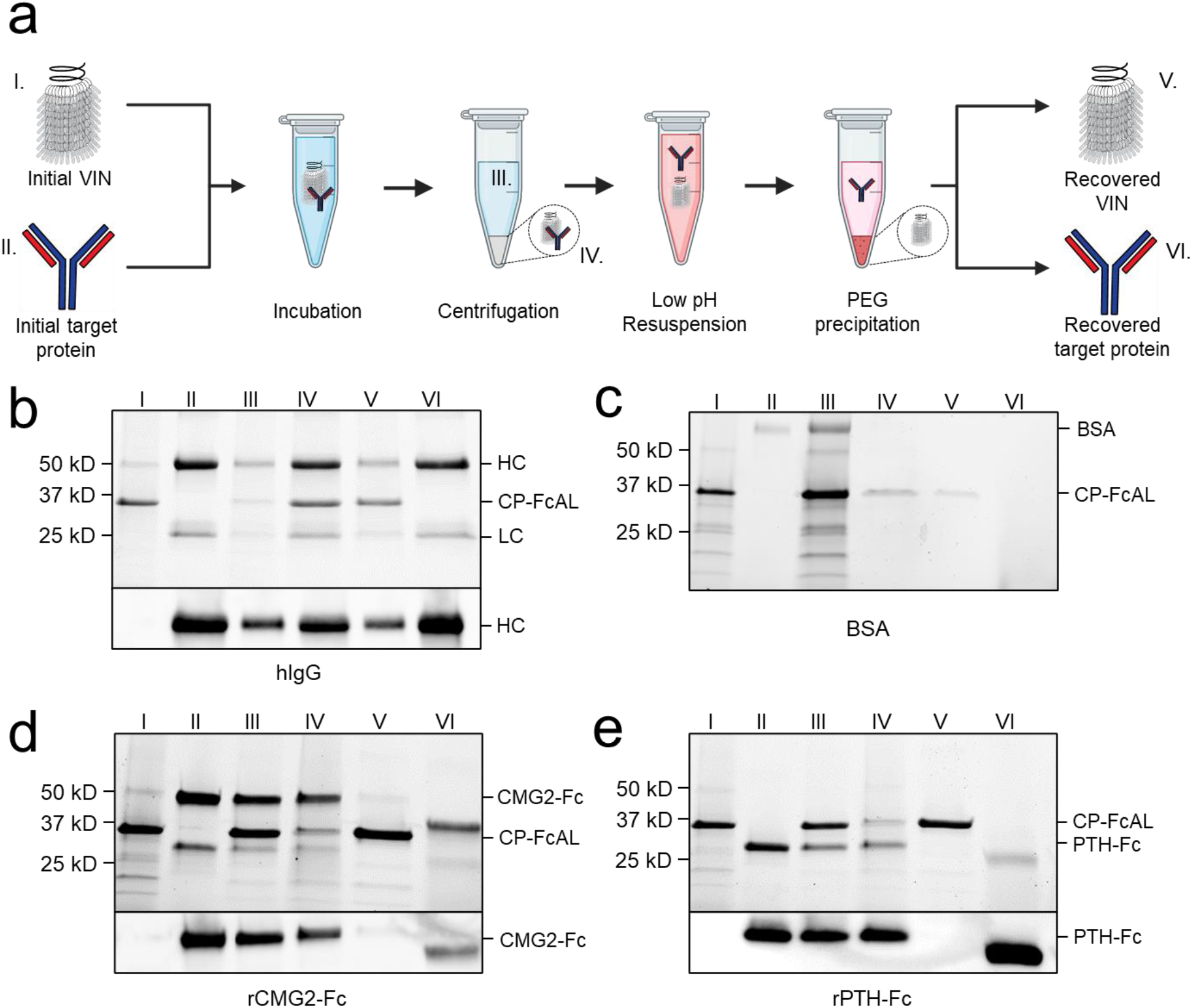
Plant virus-based immunosorbent nanoparticle (VIN)-based capture and elution of Fc-proteins from a purified solution. (a) An illustration of the VIN-based capture and elution that indicates sample points. SDS-PAGE and Western blot results of VIN-based capture and elution with pre-purified targets of (b) human immunoglobulin G (hIgG) – reduced into heavy chain (HC) and light chain (LC) constituents, (c) bovine serum albumin (BSA), (d) plant-expressed recombinant capillary morphogenesis protein Fc-fusion (rCMG2-Fc), and (e) plant-expressed recombinant parathyroid hormone Fc-fusion (rPTH-Fc). Lane definitions: I – initial VIN added; II – initial target added; III – VIN/target supernatant (loss); IV – VIN/target pellet resuspension (capture) (2x); V – recovered VIN (a,b – 2x; d,e – 4x); VI – recovered target protein (eluate) (5x).

VINs are also capable of capturing and eluting Fc-fusion proteins (Figure 2d – e). We successfully tested two pre-purified plant-expressed Fc-fusion proteins with VIN-based capture and elution: recombinant capillary morphogenesis protein Fc-fusion (rCMG2-Fc) and recombinant parathyroid hormone Fc-fusion (rPTH-Fc). We observed that the biophysical characteristics (e.g., molecular mass, Svedberg coefficient) of the domain fused to the Fc region is critical to the sedimentation step (denoted III in Figure 2a) performance.

The hIgG sedimentation conditions (12,000 x g, 10 minutes) were not adequate for rCMG2-Fc (100 kDa) and rPTH-Fc (55 kDa). We determined that 20,000 x g for 20 minutes was adequate for sedimentation of the tested Fc-fusion proteins when bound to VINs (screening data not shown). Similarly, the smaller sizes of the Fc-fusion proteins as compared to the hIgG required a higher PEG concentration (25% w/v) for the PEG-based buffer exchange step (screening data not shown). Further optimization is required to remove residual PEG in this higher concentration method, as can be observed by the PEG interference of electrophoresis (lanes VI in Figure 2d – e), although it has been shown that the presence of PEG does not impede performance of subsequent downstream processing operations including ion exchange and affinity chromatography (Roe, 2001).

### 2.3 Process characterization

We evaluated process performance from the perspective of nanomaterial stability, capture and elution functionality, and particle integrity. VINs are stable throughout the freeze-thaw process for up to 12 cycles without noticeable degradation of CP-FcAL when stored at either −20 °C or −80 °C. Long-term stability of VINs was evaluated over a series of timepoints (2 weeks, 4 weeks, 8 weeks), temperatures (−20 °C, 4 °C, 20 °C), and protease inhibitors (none, 2 mM ethylenediaminetetraacetic acid (EDTA) and 1 mM phenylmethylsulfonyl fluoride (PMSF)) (Supporting figure: Figure S7). VIN CP-FcAL were intact over the duration evaluated at −20 °C and 4 °C, while the addition of protease inhibitors was shown to prolong stability at 20 °C, with no discernable degradation observed for up to 2 weeks (data not shown).

Next, we demonstrated that the VIN functionality is retained when using samples of hIgG spiked into wild-type *N. benthamiana* plant extract (Figure 3a). Comparable performance was observed when crude *N. benthamiana* extracts of VIN were used in conjunction with antibodies spiked into crude *N. benthamiana* extracts (data not shown). Furthermore, we were able to demonstrate that the VINs recovered from a single capture and elution cycle can be reused for an additional cycle (Figure 3b). A minor fraction of hIgG was recovered with the VIN in both cycles, suggesting that hIgG recovery could be improved by optimization of the elution step (e.g., CP-FcAL binding affinity, buffer composition). The VIN recovered from the second use cycle could not be used for a third cycle with the established sedimentation conditions.

**Figure 3.**
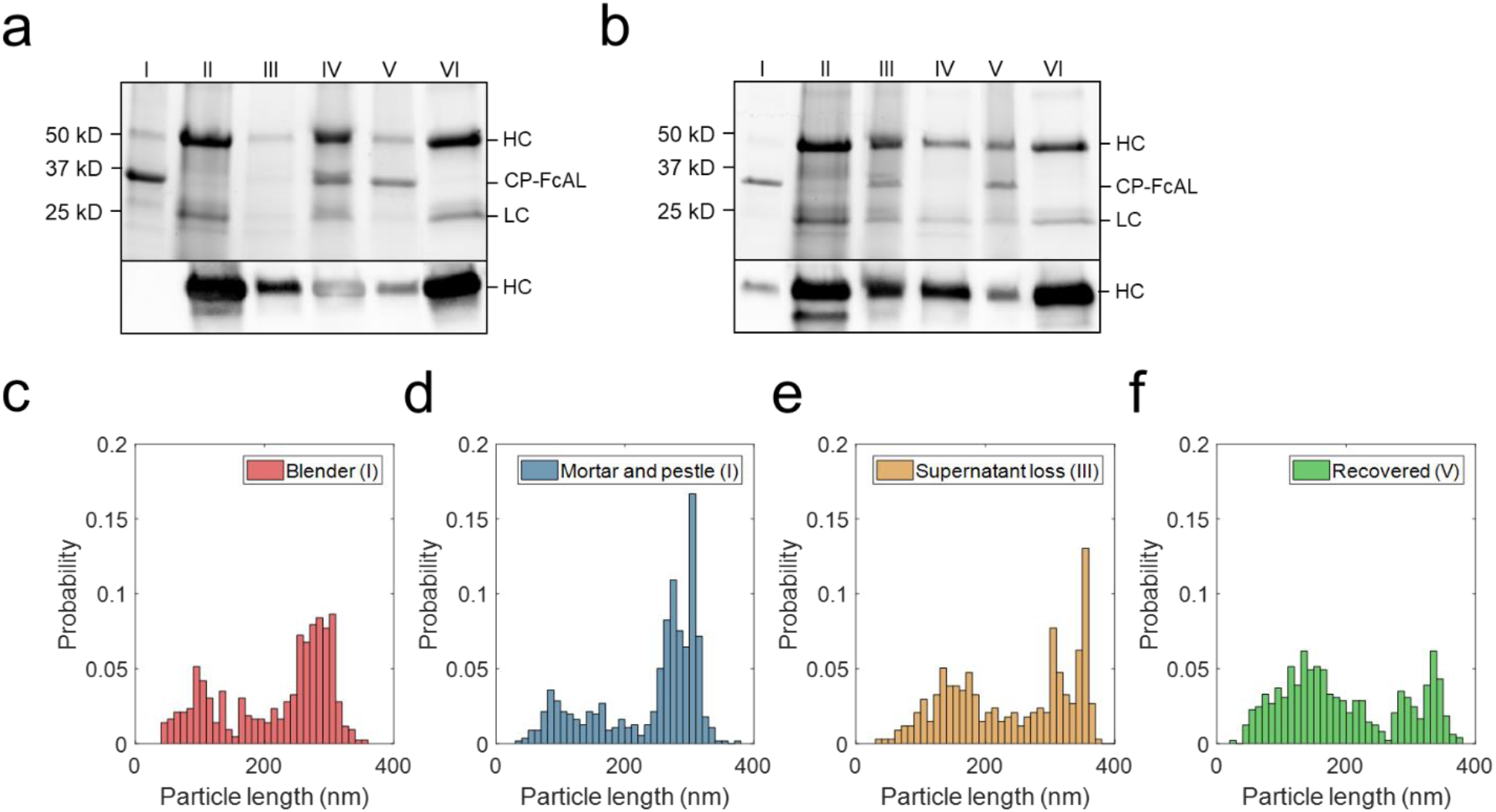
Plant virus-based immunosorbent nanoparticle (VIN)-based capture and elution of human immunoglobulin G (hIgG) from crude solution over multiple use cycles. (a) VIN-based capture and elution using a sample of hIgG in crude *N. benthamiana* plant extract, and (b) a second capture and elution cycle of the VIN recovered after the first use cycle. Lane definitions: I – initial VIN added; II – initial target added; III – VIN/target supernatant (loss); IV – VIN/target pellet resuspended (capture) (2x); V – recovered VIN (2x); VI – recovered target (eluate) (5x). Gels are marked with band heights corresponding to VIN coat protein Fc-affinity ligand fusion (CP-FcAL) and hIgG heavy chain (HC) and light chain (LC) constituents. Particle length analysis of negative stain transmission electron microscope images for (c) blender-extracted VIN, (d) liquid nitrogen-assisted mortar and pestle-extracted VIN, (e) VIN in the supernatant lost during the VIN-target complex sedimentation stage, and (f) recovered VIN post-elution. Data from parts e – f are generated using mortar and pestle extracted VIN and the naming convention I, III, and V corresponds to that established in Figure 2a.

Accordingly, VIN particle integrity was investigated to probe limitations of the sedimentation method. We first compared VINs generated by two different methods of plant extraction, a blender or liquid nitrogen-assisted mortar and pestle (Figure 3c – d). We observed a statistically significant difference in the mean particle length between the two extraction methods (p < 0.001), with blender-based extraction resulting in a shorter mean VIN length (blender: x̅ = 217 nm, σ = 84 nm, N = 428; mortar and pestle: x̅ = 239 nm, σ = 80 nm, N = 558).

We then investigated the VIN particle lengths at steps throughout the capture and elution with initial VIN generated using liquid nitrogen-assisted mortar and pestle extraction, focusing on the VINs lost in the supernatant during the VIN-hIgG complex sedimentation step (Figure 3e) and the final recovered VINs (Figure 3f), denoted III and IV in Figure 2a, respectively. There is an observed statistically significant difference in the mean particle length for the initial VINs, VINs lost in the supernatant (x̅ = 241 nm, σ = 94 nm, N = 337), and final recovered VINs (x̅ = 196 nm, σ = 93 nm, N = 486) (I & III, p = 0.039; I & IV, p < 0.001; III & IV, p < 0.001).

### 2.4 Process development

The appreciable level of impurities present in the purified VIN solutions, as well as an interest in improving scalability of the processing by removing the chloroform-based liquid-liquid extraction step, motivated an investigation into process development of the VIN purification. We performed a 2-factor 2-level process optimization of the extraction step (buffer composition – 50 mM sodium acetate 86 mM NaCl pH 5.0, 100 mM potassium phosphate pH 7.0; protease inhibitors – none, 2 mM EDTA + 1 mM PMSF) followed by an addition of a heat hold step that was investigated with a temperature screening (30 – 70 °C) after each processing operation (Figure 4a).

**Figure 4.**
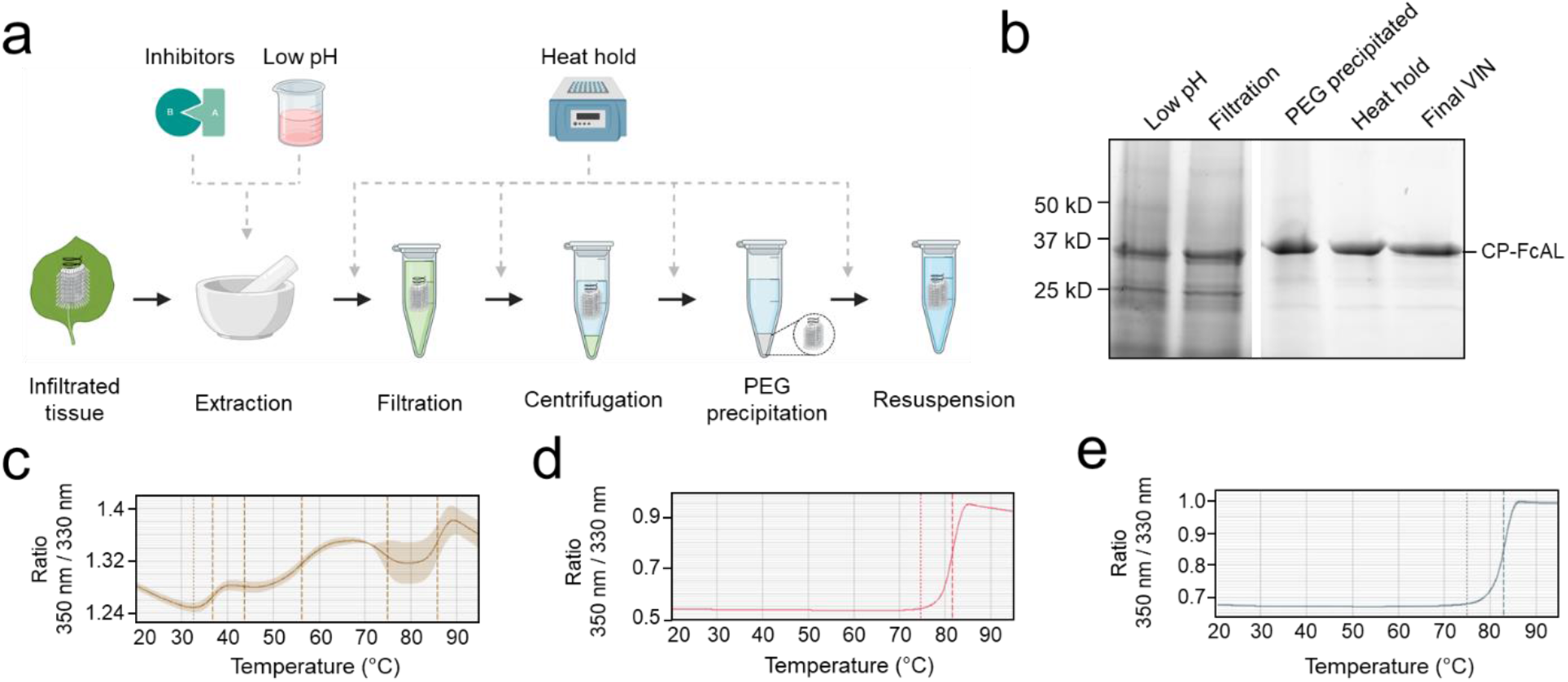
A summary of the plant virus-based immunosorbent nanoparticle (VIN) preparation process improvement conducted in this study. (a) An illustration of the VIN preparation stages and the experimental design tested, and (b) SDS-PAGE results of the optimized VIN preparation scheme shown marked at the band height corresponding to VIN coat protein Fc-affinity ligand fusion (CP-FcAL). Nano differential scanning fluorimetry assessment of protein thermostability is shown for (c) crude extract at pH 5, (d) VIN prepared according the optimized scheme, and (e) VIN prepared according to the baseline purification scheme, using the intrinsic tryptophan and tyrosine residue fluorescence at 350 nm and 330 nm.

Results of the process optimization conditions tested, including verification of bind-and-elute functionality of VIN produced from the different purification schemes, can be found in Supporting figures: Figure S8, Figure S9, and Figure S10.

The chloroform-based liquid-liquid extraction step was removed from the processing scheme with comparable or improved VIN recovery and purity upon inclusion of a low pH extraction and 60 °C heat hold post-PEG precipitation (Figure 4b). Interestingly, the addition of protease inhibitors to the extraction buffer reduced VIN recovery and increased the presence of what appears to be degradation products. Similarly distinct from wild-type virion processing, the heat hold resulted in significant VIN loss when introduced at processing steps prior to PEG precipitation. This behavior can be attributed to the FcAL presentation, as wild-type tobacco mosaic (wt-TMV) is routinely processed with early-stage processing heat holds (Smith *et al.*, 2006).

Nano differential scanning fluorimetry results (Figure 4c – e) indicate that the VINs prepared according to either protocol detailed in this study exhibit a melting temperature of ~82 °C, supporting that the improved protocol does not introduce discernible differences in VIN CP-FcAL stability. There are multiple distinct conformational shifts within the crude solution consistent with the heterogeneity of solution.

### 2.5 Magnetic separation with VIN

Figure 5a illustrates the basic concept and utility of the VIN-coupled magnetic particles (VIN-MPs) generated in this study and Figure 5b – c displays TEM images of the intact VIN-MPs complex. A magnetic separation-based capture and elution method was developed with hIgG spiked into phosphate buffered saline (PBS) (not shown) and crude *N. benthamiana* plant extract that confirms the VIN immunosorbent functionality in this novel configuration (Figure 5d). A faint presence in the elution that may indicate that some minor amount of VIN is recovered in addition to the target hIgG.

**Figure 5.**
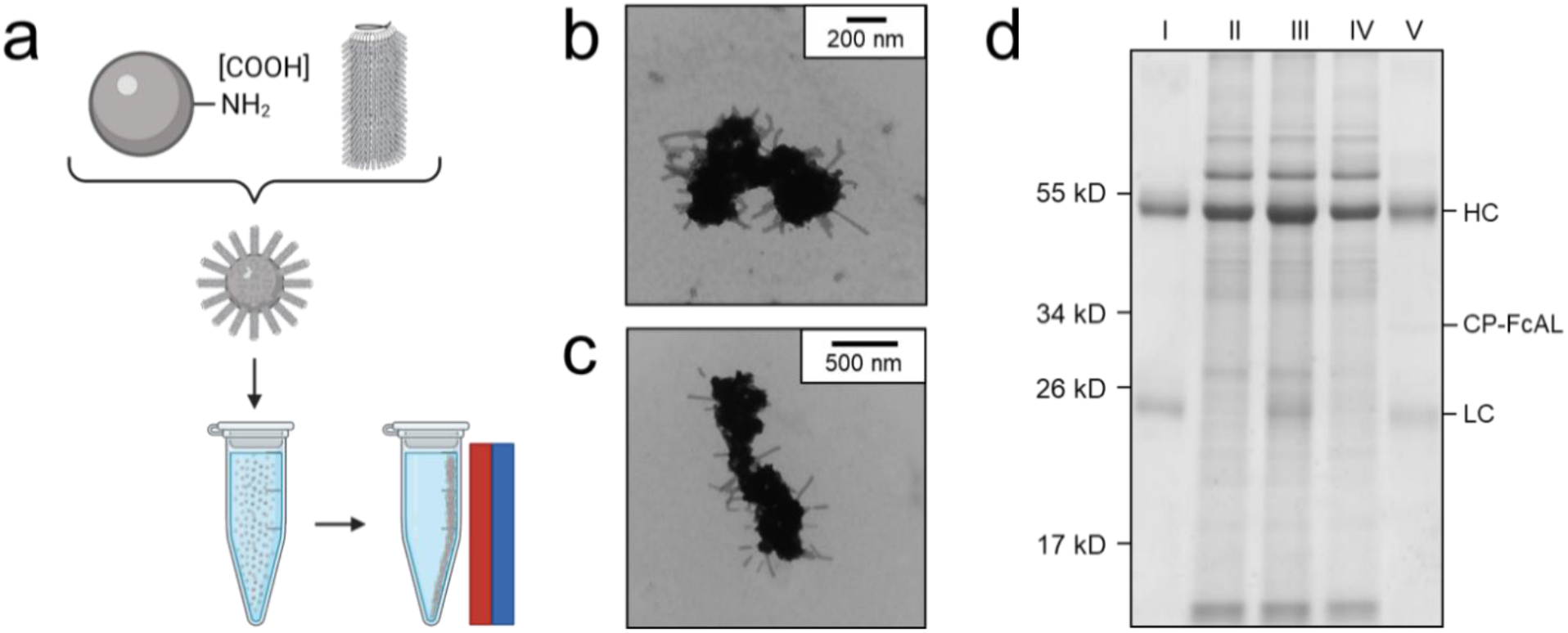
Production of plant virus-based immunosorbent nanoparticles coupled with magnetic particles (VIN-MPs). (a) An illustrative depiction of VIN-MPs. (b, c) Negative stain transmission electron microscope images of VIN-MPs generated using amine-terminated magnetic particles and 5% glutaraldehyde (VIN-MP-NGs). (d) Reducing condition SDS-PAGE of human immunoglobulin G (hIgG) bind-and-elute using VIN-MP-NGs and magnetic separation. Gels are marked with band heights corresponding to VIN coat protein Fc-affinity ligand fusion (CP-FcAL) and hIgG heavy chain (HC) and light chain (LC) constituents. Lane definitions: I – initial hIgG added; II – *N. benthamiana* plant extract; III – hIgG spiked into *N. benthamiana plant* extract; IV – non-bound supernatant after magnetic separation with VIN-MPs; V – hIgG elution from VIN-MPs.

The VIN-MP were either coupled with amine-terminated (VIN-MP-N) or carboxyl-terminated (VIN-MP-C) superparamagnetic particles. The VIN-MP-N coupled at significantly higher densities (> 0.2 mg VIN/mg MP) than the VIN-MP-C (≤ 0.1 mg VIN/mg MP) and resulted in higher hIgG capture. Therefore, VIN-MP-N were selected as the basis for additional study.

Two different coupling agents were tested: glutaraldehyde (VIN-MP-NG) and 1-Ethyl-3-(3-dimethylaminopropyl)carbodiimide (EDC) (VIN-MP-NE). Furthermore, we screened a range of EDC concentrations for use in the VIN-MP-NE synthesis reaction. We observed that the coupling density of VINs to MPs could be tuned with the concentration of EDC used in the covalent coupling reaction (Figure 6a). We also observed that the VIN concentration in the reaction medium could be used to tune coupling density (data not shown).

**Figure 6.**
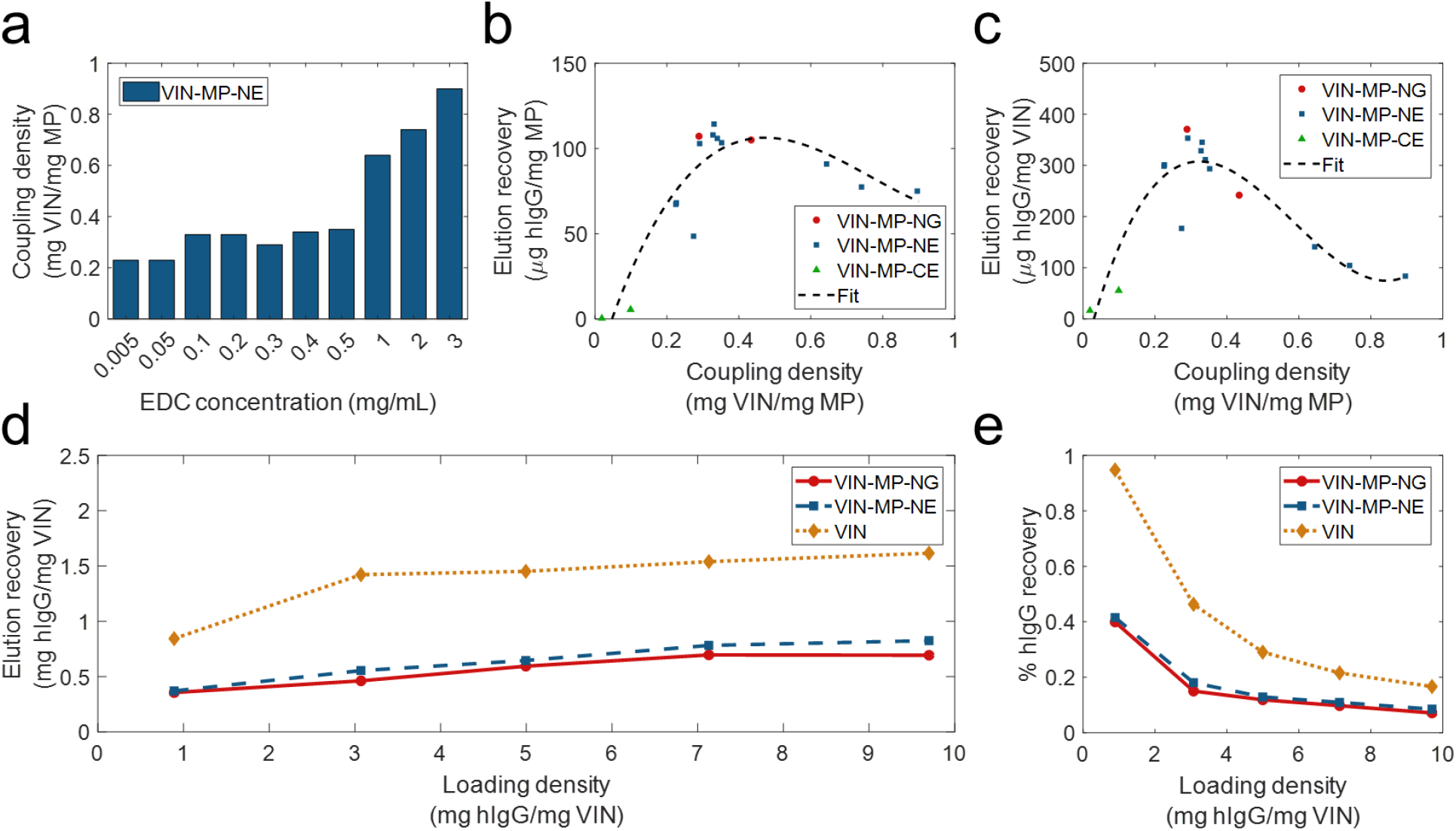
Screening and evaluation of plant virus-based immunosorbent nanoparticle-coupled magnetic particle (VIN-MP) coupling density and human immunoglobulin G (hIgG) elution. (a) 1-Ethyl-3-(3-dimethylaminopropyl)carbodiimide (EDC) concentration used during coupling and resultant coupling density for amine-terminated magnetic particles. VIN-MP coupling density and hIgG elution recovery for both amine- and carboxyl-terminated magnetic particles (b) per MP mass basis, and (c) per VIN mass basis. Fits are generated as 3^rd^ order polynomials. Elution recoveries of hIgG from PBS are shown over a range of hIgG loading densities for amine-terminated MP (VIN-MP-N) with 5% glutaraldehyde coupling (VIN-MP-NG), 0.5 mg/mL EDC coupling (VIN-MP-NE), and uncoupled VIN in free suspension (d) per VIN mass basis, and (e) as an extent of the loading density and of hIgG from PBS.

We observed a non-monotonic relationship between coupling density and hIgG binding/elution load per mass of MP (Figure 6b). An optimal coupling density of 0.3 – 0.4 mg VIN/mg MP was identified. The hIgG binding/elution load per mass of VIN was consistent at approximately 0.3 mg hIgG/mg VIN below a coupling density of 0.4 mg VIN/mg MP, above which a negative correlation between coupling density and hIgG binding/elution load per mass of VIN is observed (Figure 6c). This suggests that higher coupling densities may provide less available hIgG binding sites due to steric hinderances or electrostatic interactions.

A comparison of VIN capture and elution performance with hIgG in PBS is shown in Figure 6d-e for magnetic separation using VIN-MP and sedimentation separation using uncoupled VIN over a range of hIgG loading densities. The VIN-MP hIgG elution recovery normalized by mass of VIN is approximately 35-55% of that of the uncoupled VIN.

Sedimentation and magnetic separation-based operations were further compared in purification of plant-expressed monoclonal antibody (mAb) A20 from crude *N. benthamiana* plant extract (Figure 7). Non-reducing condition SDS-PAGE results indicate that fully assembled mAb A20 is produced and recovered by VIN in elution. The relative extents of hIgG elution recovery for uncoupled VIN and VIN-MP are consistent with the hIgG in PBS results.

**Figure 7.**
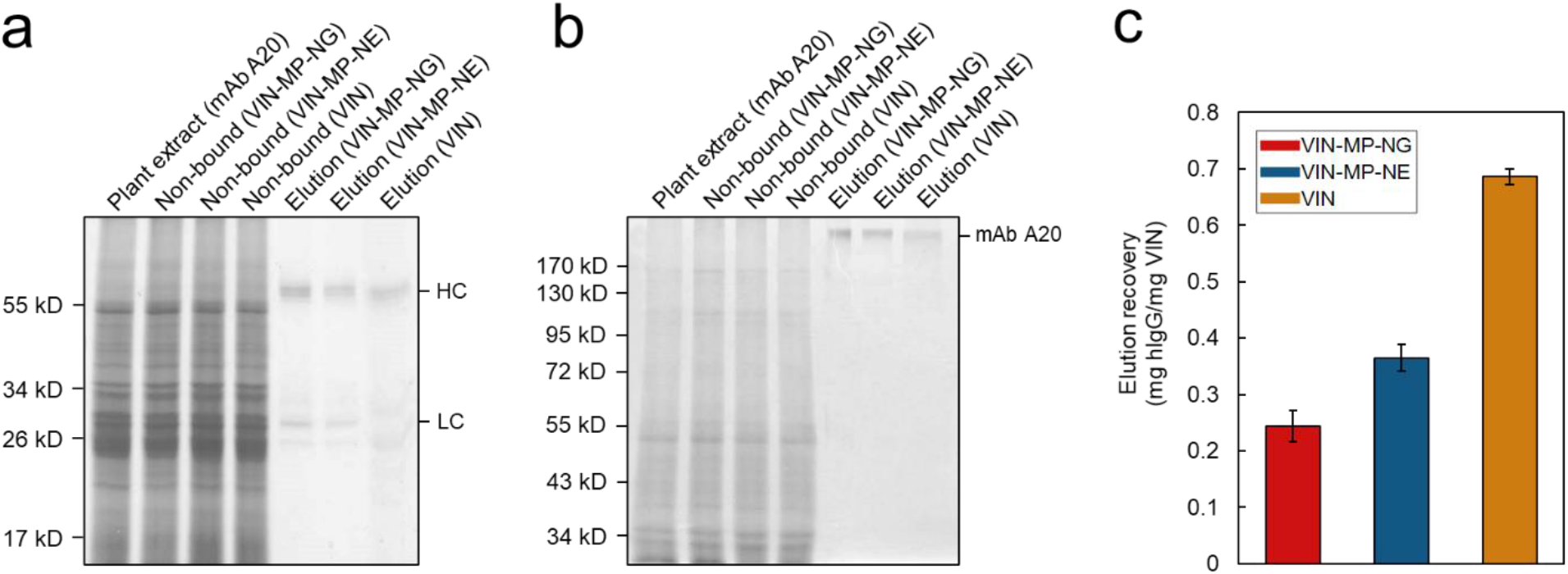
Capture and elution of plant-expressed monoclonal antibody (mAb) A20 by sedimentation with uncoupled plant virus-based immunosorbent nanoparticle (VIN) and magnetic separation with VIN-coupled magnetic particle (VIN-MP). (a) Reducing and, (b) non-reducing SDS-PAGE results of the non-bound supernatant and elution recovery of plant-expressed mAb A20 for VIN-MP with 5% glutaraldehyde coupling (VIN-MP-NG), 0.5 mg/mL 1-Ethyl-3-(3-dimethylaminopropyl)carbodiimide coupling (VIN-MP-NE), and uncoupled VIN in free suspension. Gels are marked with band heights corresponding to mAb A20 heavy chain (HC) and light chain (LC) constituents (reduced) or to dimerized mAb A20 (non-reduced). (c) Elution recovery of plant-expressed mAb A20 for VIN-MP-NG, VIN-MP-NE, and uncoupled VIN in free suspension. Error bars represent a single standard deviation of technical triplicate measurements.

## 3. Discussion

### 3.1 Biotic purification technologies

Virus-based nanomaterials present promising characteristics as an alternative technological platform to traditional chemical methods of biopharmaceutical purification in their inexpensive and scalable production coupled with their high replication fidelity, biophysical properties, stability, and accessible modifications leading to wide-ranging functionality. There are safety and regulations concerns of commercializing self-replicating technology, but this barrier has been addressed by researchers by removing the requisite replication machinery, as is done to generate virus-like particle technology (Marsian and Lomonossoff, 2016; Zeltins, 2013), using a plant virus to avoid human infection (Balke and Zeltins, 2019), designing the virus-based nanomaterial to rapidly shed the transgenic gene inserts (Torti *et al.*, 2021), and/or inactivating the virus (Koudelka *et al.*, 2015). As these methodologies are well established, we do not address virus nanoparticle containment strategies within the scope of VIN process development.

A range of biotic technologies beyond virus-based nanomaterials have been developed and studied for biopharmaceutical purification (Dias and Roque, 2016; Mahmoodi *et al.*, 2019). These can be classified by utility as fusion tags (e.g., inteins (Belfort *et al.*, 1999), carbohydrate binding modules (Shoseyov *et al.*, 2006)), thermo-responsive biopolymers (e.g., elastin-like polypeptides (Sheth, Bhut, *et al.*, 2014)), and hydrophobic nanoparticles (e.g., polyhydroxyalkanoates (Banki *et al.*, 2005), oleosins (Bhatla *et al.*, 2010), hydrophobins (Jugler *et al.*, 2020)). Fusion tags have by and large been the most widely adopted biotic purification technology for the accessibility they present to early-stage research labs. However, they are limited as a platform technology by the influence of product-specific characteristics, complications of tag cleavage, and generally unfavorable commercial-scale economics (Fc-fusion tags being a notable exception to most of these limitations) (Bell *et al.*, 2013). There has been some adoption and maturation of the other biotic technologies that overcome these limitations, with an observed emphasis on elastin-like polypeptides (ELPs) (Sheth, Bhut, *et al.*, 2014; Sheth, Jin, *et al.*, 2014) and oleosins (McLean *et al.*, 2012). VINs represent another contender within this group of technology, albeit at a more nascent stage of development. Advantages of VIN technology include the simplicity of production (ELP: culture-based system; VIN: plant-based system) and within that the projected high yield per hectare (oleosin: < 1 kg/hectare; VIN: 200+ kg per hectare) (Werner *et al.*, 2006) that position VINs as inexpensive purification reagents.

Mechanical transmission-based production of VINs, as we have demonstrated, could be considered for economical manufacturing for its enhanced simplicity over agrobacterium-based methods and its reliability and stability over transcript-based methods. A main barrier to this strategy is the variability and escape of the VIN functionality over the course of multiple virion replication and plant passage cycles. One strategy to alleviate these concerns would be to embed selective pressures into the processing procedure, although this would introduce yet unsolved barriers within quality assurance and quality control that would hinder commercialization. Studies have shown the use of selective pressures during production can cause viruses to sacrifice reproductive fitness for selected characteristics (e.g., thermal and structural stability) (Dessau *et al.*, 2012) and that genetic stability of virus-based nanomaterials can be achieved (Le Nouën *et al.*, 2017).

### 3.2 Affinity sedimentation processing

These technologies have been applied in a diverse range of processing strategies including liquid chromatography, inverse transition cycling, aqueous two-phase partitioning, and affinity precipitation. The uncoupled VIN application methodology presented here is based on what we are terming as pseudo-secondary effect affinity sedimentation, in which the affinity interaction and sedimentation mechanisms are partially coupled with a dependence of the sedimentation on affinity interaction (i.e., VIN-target protein complex characteristics influence sedimentation velocity). We derive this terminology from affinity precipitation processing which has distinguished methodologies as either primary effect (coupled affinity/precipitation) or secondary effect (independent affinity/precipitation) (Hilbrig and Freitag, 2003).

We performed initial work to identify centrifugation conditions as a function of target protein characteristics and loading (e.g., antibody versus Fc-fusion protein), but it may be valuable in future works to develop a model to understand this relationship more deeply between the VIN-target protein complex morphology and sedimentation velocity. It was qualitatively observed that Fc-fusion protein recovery was lower than that of hIgG regardless of centrifugation conditions. We hypothesize that lower binding affinities and differing biophysical characteristics of the VIN-Fc-fusion protein complex are contributing to these observed differences. However, future work is required to elucidate these underlying mechanisms.

Affinity sedimentation shares several characteristics with affinity precipitation, which provides potential benefits of low cost and buffer usage (Hilbrig and Freitag, 2003), ability to achieve high concentration factors (Low *et al.*, 2007), high throughput (Shukla and Thömmes, 2010), and minimal concerns of fouling at the expense of higher recovery and selectivity generally achieved by a chromatographic counterpart (Mondal *et al.*, 2006), which can consist of as much as 50% of the total pharmaceutical manufacturing costs (Kelley, 2009). Affinity sedimentation and precipitation methods also exhibit generally higher tolerance to variation in feed streams, as we have shown with VINs through processing of crude plant extract, making them well-suited to early-stage downstream processing.

A semi-quantitative comparison of hIgG binding capacity between the VINs prepared at two levels of purity (data not shown) indicates that there may be a minor reduction in binding capacity between these conditions, likely associated with non-specific blocking by various expected plant host impurities including proteins, salts, polysaccharides, and phenolics (Dixon *et al.*, 2018). However, there was no discernable impact of the solution type (crude extract or purified solution) on reusability of the VINs for an additional cycle of capture and elution.

While protein solids, such as those formed in precipitation and sedimentation, have been shown to be stable in long term storage (Harrison *et al.*, 2015), here we show that sedimentation impacts VIN structural integrity and presumably contributes to the unstable performance over multiple reuse cycles. From these results, one may infer the importance of fully intact VIN on sedimentation characteristics and thus centrifugal recovery. The mechanism of particle breakage is suspected to be mechanically induced from during resuspension and not a direct result of the protein pellet formation from sedimentation. The mechanical properties of viruses have been studied extensively using computational and physical methods (Buzón *et al.*, 2020; Mateu, 2012). TMV, in the same genus as the TVCV used for the VIN in this study, has been attributed a Young’s modulus of 6 ± 3 GPa (Schmatulla *et al.*, 2007), although mixed reports suggest the value could be lower (Falvo *et al.*, 1997). Furthermore, there are reports on icosahedral virions that demonstrate minor changes in coat protein composition resulting in significant modulation of mechanical properties (Medrano *et al.*, 2019).

Thus, we hypothesize that the presumably stiff TVCV rod-like particle basis combined with increased drag by the loose affinity ligand display sheath (using the largest genetically inserted virion coat protein presentation to date) surrounding the VIN enhances its vulnerability to applied shear stress during pellet resuspension via repeated pipette tip aspiration. It may be that a smaller affinity ligand presentation, such as an affibody (Frejd and Kim, 2017) or synthetic peptide (Lund *et al.*, 2012), would decrease particle breakage (but also necessitate more aggressive centrifugation conditions with the smaller virion size). Gentler resuspension methods with lower shear would also be worth investigating to decrease breakage. Similar shear sensitivity in pipette-based resuspension has been shown for larger biomolecule systems such as with cell-based pelleting, wherein higher pellet compaction and tip velocities were shown to result in cell losses (Delahaye *et al.*, 2015). Pellets formed during VIN processing and use were highly compacted and required considerable pipetting for complete resuspension, suggesting that pipette-based resuspension could have also played a role of particle degradation in this system. Gentler resuspension techniques should be explored in the future to improve particle integrity during operation.

In this study, we investigated stability of the VINs over long term storage, multiple freeze-thaw cycles, and at elevated temperatures, primarily focusing on coat protein fusion primary structure. Additional stability concerns include impact to protein secondary structure, virion particle structure, and nucleic acid integrity. Exposure to multiple freeze-thaw cycles has been shown to degrade virion nucleic acid and infectivity (Krajden *et al.*, 1999). Thermostability was confirmed at the level of secondary protein structure via performance check of bind-and-elute functionality, which led to the integration of a high heat hold into the VIN purification scheme.

### 3.3 VIN process development

The process development investigation presented here provides insights on the differences between wild-type plant virions and protein display plant virions, which is relevant technology for a host of biomedical applications. The improved process can serve as a roadmap for future virus-based nanomaterial purification. We observed that the protein presentation confers additional processing sensitivities to the virion (which is otherwise described as a glassy surface), likely due to the interactions between the presented ligand and *N. benthamiana* plant host cell impurities, noticeable in the heat hold of crude solution and the inclusion of protease inhibitors to extraction.

We observed that the two effective unit procedure modifications, low pH extraction and heat hold post-PEG precipitation, possess low orthogonality in impurity clearance mechanisms for the starting stream used although there is still discernable benefit in combining the methods as observed by the improvement in SDS-PAGE band purity.

The process development in this work focused on addition and removal of unit procedures at a high-level to inform process design. There is value in future research performing parameter optimization with an emphasis on maximizing recovery and purity with a fixed purification scheme.

### 3.4 Magnetic separation

The VIN-MP results presented in this study, representing the first virus-based nanomaterial system coupled with MPs for protein purification, reflect a greater than 25x increase in binding capacity compared to current industry standards for affinity protein capture with magnetic particles – Pierce^TM^ Protein A Magnetic Beads (≥ 40 μg rabbit IgG/mg MP) (ThermoFisher Scientific), SureBeads^TM^ Protein A Magnetic Beads (≥ 6 μg IgG/mg MP) (Bio-Rad Laboratories), VIN-MP (> 1,000 μg hIgG/mg MP).

These exciting results provide stark evidence for the sensitivity-enhancing properties of virus-based nanomaterials and their usefulness as ligand scaffolding in biotechnological applications.

Furthermore, the VIN-MP system served to decouple the affinity and separation mechanisms of processing, as compared to the partially coupled behavior in VIN sedimentation operation, thereby increasing the process robustness to changes in the sample solution, including the diversity and concentration of the target protein. Additional work is required to experimentally assess this capability in a larger set of processing conditions. Investigation of the reusability of VIN-MP and for magnetic separation and particle integrity over operation is also of importance for future testing. Preliminary results suggest there may be a minor presence of VIN in the eluate. Two possible means of explaining this observation are proteolytic cleavage along a covalently bonded FcAL resulting in detachment of the VIN from MP or minor particle breakage from resuspension of the VIN-MP after magnetic separation resulting in the presence of VIN fragments.

The uncoupled VIN sedimentation operation demonstrated 2-3 times higher capture capacity per VIN mass than the VIN-MP system. The uncoupled VIN operation also resulted in hIgG recoveries as high as 95% of the feed, whereas VIN-MP operation was maximal at 42% recovery of the feed hIgG. It will be important to identify the cause for the lower recovery in future development by additional screening of lower hIgG concentrations and optimization of the capture and elution methodology.

Higher capture capacity may be particularly advantageous for large-scale protein purification – an area for which affinity precipitation (Swartz *et al.*, 2018) and magnetic separation (Schwaminger *et al.*, 2019) are receiving growing interest. However, given the nature of VINs as inexpensive and simply produced reagents, the value of maximizing Fc-protein binding per VIN in a small-scale commercial application (as is the current niche of magnetic particle purification) is most likely weighted less than factors such as Fc-protein recovery, process duration, labor time, amenability to automation, and equipment costs.

For perspective on these other factors, consider that our recently published study on evaluating the costs of the affinity capture step of mAb purification (McNulty *et al.*, 2021) yielded results that the unit costs of magnetic separation (modeled using an industry standard technology with a comparable capture and elution protocol) were lower than uncoupled VIN sedimentation in process duration (73% reduction), labor time (30% reduction), and equipment mass (71% reduction) when processing a single lab-scale sample (2 mL volume tube). These results support the favorable position of VIN-MP in comparison to uncoupled VIN sedimentation for lab-scale applications.

### 3.5 Summary and future directions

Virus-based nanomaterials provide a highly diverse and tunable technology that can be adapted to overcome the limitations of the application methodology. For example, the length of a rod-like plant virus such as the one used in this study, TVCV, is proportional to the length of the viral genomic information and, as such, the length can be modulated through the addition or subtraction of genomic information (e.g., addition of non-functional genomic information can be used to increase virion particle length) (Saunders and Lomonossoff, 2017). Increasing VIN length in this manner is one approach to investigate for increasing the binding site occupation in the VIN-MP system. Other techniques useful for VIN performance optimization include density modulation of the protein display (Cruz *et al.*, 1996) and a multi-ligand protein display (Werner *et al.*, 2006). Last not least, use of affinity ligands other than Protein A domains, in particular, affibodies evolved from Protein A/Z domains (Nord *et al.*, 1997), should expand the usability of the technology beyond monoclonal antibody binding/capture (Ståhl *et al.*, 2017).

In this study, we have presented the development of a virus-based nanomaterial to serve as a protein purification reagent, characterized performance using a pseudo-secondary effect affinity sedimentation bind-and-elute protocol, expanded functionality to Fc-fusion proteins, identified limitations of the technology operated in that procedure, and developed a magnetic particle coupled system for magnetic separation to improve processing. This provides further evidence supporting virus-based nanomaterials as simple and inexpensive reagents for protein purification and suggests a path forward for technological development.

## 4. Experimental procedures

### 4.1 Gene constructs

The viral expression vector used in this study is based on previously reported TVCV-based vectors(Werner *et al.*, 2006). The viral expression vector used here (pICH25892; plasmid kindly provided by Nomad Biosciences GmbH) is an assembly of the previously reported 5’ provector containing the TVCV coat protein (minus the stop codon) fused to a C-terminal glycine-rich flexible linker (pICH20701) and the 3’ provector containing the D and E antibody-binding domains from *S. aureus* protein A with short flanking sequences (amino acids 29 – 161; GenBank accession no. J01786) (pICH21767).

### 4.2 Production of VIN

VINs were primarily produced via whole plant agroinfiltration using *A. tumefaciens* containing viral expression vector pICH25892 according to a previously reported method with minor modifications (Xiong *et al.*, 2018). A final cell density of OD_600_ = 0.2 was used for agroinfiltration. Post-infiltration plants were cultivated at 60% relative humidity with a 16-hour photoperiod, 23 °C/20 °C temperature regime, and a photosynthetic photon flux density of 425 μmol/(m^2^∙s) derived from a combination of high-pressure sodium, high-pressure metal halide, and incandescent lights for a duration of 6-12 days post-inoculation.

VINs were also produced via direct mechanical transmission of intact VINs. A total volume of 300 μL of purified VIN solution (~0.1 mg/mL) was applied per plant in aliquots of 100 μL for each of three middling leaves. An abrasive powder (Celite) was lightly sprinkled on each leaf and each leaf was gently rubbed by hand. The surfaces of the leaves were rinsed with water at 20 minutes post-inoculation to remove excess inoculation reagents.

### 4.3 Purification of VIN

VIN-expressing *N. benthamiana* leaf tissue was stored at −80 °C after harvest and processed with minor modifications to a previously reported protocol (Werner *et al.*, 2006). A single round of PEG-assisted precipitation step was performed rather than two. Extraction was performed using either a blender (NutriBullet; NutriBullet, LLC, Pacoima, CA) or liquid nitrogen-assisted mortar and pestle with 0.1 M potassium phosphate pH 7.0 extraction buffer at a 3:1 buffer volume to biomass weight extraction ratio. In the case of the mortar and pestle method, the homogenized leaf powder was mixed with the buffer and nutated for 30 minutes at 4 °C for extraction.

### 4.4 Binding and elution of Fc-proteins

Binding and elution of Fc-proteins was performed according to a previously reported protocol (Werner *et al.*, 2006). Four different target Fc-proteins were used in this study: hIgG (Sigma-Aldrich, St. Louis, MO), plant-expressed rCMG2-Fc) (Xiong *et al.*, 2019), plant-expressed rPTH-Fc (unpublished data), and using the magnICON^®^ system (Giritch *et al.*, 2006), plant-expressed mAb A20 (subclass IgG1) (Bendandi *et al.*, 2010; Whaley *et al.*, 2012). Development of target protein-specific modifications to the method are detailed in results.

### 4.5 Coupling VIN with magnetic particles

VIN-MP were generated by covalent attachment of VIN to primary amine-terminated superparamagnetic iron oxide particles (Product No. I7643, Sigma-Aldrich, St. Louis, MO, USA), VIN-MP-N, or carboxyl-terminated superparamagnetic iron oxide particles (Product No. I7518, Sigma-Aldrich, St. Louis, MO, USA), VIN-MP-C, both of approximately 1 μm size, was performed according to the methods detailed in the product data sheets using ~2 mL total reaction volumes. VIN stock solutions at ~3-6 mg/mL concentration in 10 mM potassium phosphate buffer pH 7.0 were used in coupling. Coupling efficiency was measured using A280 values for amine-terminated particles and Bradford assay soluble protein values for carboxyl-terminated particles.

### 4.6 Binding and elution of Fc-proteins with VIN-magnetic particles

A VIN-MP solution was prepared by resuspending 2.5 mg of VIN-MP in 0.5 ml of 0.1 M sodium phosphate buffer pH 8.0 binding buffer. Further, the particles were magnetically separated; the supernatant was aspirated and discarded (repeated three times).

Crude protein extracts from *N. benthamiana* leaves (leaf juice press extraction followed by microfiltration with filter paper) in binding buffer were spiked with various amounts of hIgG. A volume of 100 μL hIgG-containing crude extract was added to the VIN-MP solution. The mixture was briefly vortexed and then incubated nutating at 4 °C for 30 minutes. The incubated solution was then magnetically separated and washed three times with binding buffer. The binding buffer was removed after wash and 50 μl of 0.2 M glycine buffer pH 2.5 elution buffer was added. The solution was briefly vortexed to resuspend magnetic particles and further incubated nutating at 4 °C for 5 min. Particles were again magnetically separated and the supernatant was collected as the eluate. The eluate was then pH neutralized with 13 μl of 1.5 M Tris-HCl buffer pH 8.8. The elution and neutralization steps were repeated three times and pooled together.

### 4.7 Protein analysis

Protein concentration was measured using Bradford and Pierce Modified Lowry assays.

Sample protein compositions were analyzed by SDS-PAGE and Western blot. SDS-PAGE samples were loaded using constant volume (30 μL). Western blot analysis was performed using a primary antibody of rabbit anti-protein A (1:25,000 dilution) (Sigma-Aldrich, St. Louis, MO) and a secondary antibody of goat anti-rabbit IgG-HRP (1:3,000 dilution) (Southern Biotech, Birmingham, AL) for detection of VIN CP-FcAL. A secondary antibody of goat anti-human IgG-HRP (1:2,500 dilution) was used to detect human IgG, rCMG2-Fc, and rPTH-Fc.

Dot blots were performed using 5 μL liquid samples and 0.45 μm nitrocellulose membrane. The positive control consisted of 100 – 500 ng recombinant Protein A (ThermoFisher Scientific, Santa Clara, CA). The negative control consisted of ~2 μg wt-TMV from purified *N. benthamiana* solution. VIN samples consisted of ~100 μg of VIN from purified *N. benthamiana* solution based on total soluble protein assay results. Several secondary antibody conditions were used: rat anti-mouse IgG-HRP (1:1,000 dilution) (ThermoFisher Scientific), rabbit anti-goat IgG-HRP (1:3,000 dilution) (Invitrogen, Carlsbad, CA), and goat anti-human IgG-HRP (1:3,000 dilution).

### 4.8 Electron microscopy

Carbon film on 300 mesh copper discs (Ted Pella, Redding, CA, USA) were prepared for increased hydrophilicity by glow discharge at 30 mA for 30 seconds on a glass slide. 5 μL liquid VIN solution samples were loaded onto the prepared disc, incubated 30 seconds, and then blotted with filter paper. Negative stain was applied in five sequential rounds of 5 μL uranyl sulfate loading, 30 second incubation, and filter paper blotting. TEM was performed using a JEM-1230 transmission electron microscope (JEOL, Peabody, MA, USA).

The lengths of VIN particles imaged by TEM were manually measured using straight line analysis with ImageJ (National Institutes of Health, Bethesda, MD, USA). Statistically significant differences in mean particle lengths of different VIN solutions were determined by equal variance two sample t-test (α = 0.05). The equal variance assumption was evaluated by two sample F-test (α = 0.05).

## 5. Author contributions

MJM, AS, AG, YG, KAM designed and executed the experiments. All authors were involved in analysis and interpretation of data. MJM wrote the initial manuscript draft. KAM edited the manuscript draft. AG, YG, SN, KAM supervised the project.

## 6. Acknowledgements

This material is based upon work supported by NASA under grant or cooperative agreement award number NNX17AJ31G. This work was also supported by a NASA Space Technology Research Fellowship (NASA grant number 80NSSC18K1157). This work is supported by the Translational Research Institute through NASA Cooperative Agreement NNX16AO69A. Any opinions, findings, and conclusions or recommendations expressed in this material are those of the author(s) and do not necessarily reflect the views of the National Aeronautics and Space Administration (NASA) or the Translational Research Institute for Space Health (TRISH).

We thank Dr. Gerd Hause (Martin Luther University of Halle-Wittenberg, Halle, Germany) for his help with electron microscopy. Illustrations were created using Biorender.com.

## 8. Short legends for Supporting Information

**Figure S1.** Agrobacterium tumefaciens T-DNA vector constructs

**Table S1.** Primers used for verifying vector construct DNA sequence

**Figure S2.** Gel electrophoresis results of PCR-based sequence verification

**Figure S3.** Amino acid analysis coverage

**Figure S4.** Dot blot of immunosorbence for various antibodies

**Figure S5.** Capture and elution of antibodies with mechanical transmission-generated particles

**Figure S6.** Capture and elution using high sedimentation rate

**Figure S7.** Freeze-thaw cycle and long-term stability

**Figure S8.** Process development observations

**Figure S9.** Thermostability assessment results

**Figure S10.** Performance verification after process development

## 10. Tables

No tables to present in this study.

