## Supporting Information for "Affinity sedimentation and magnetic separation with plant-made immunosorbent nanoparticles for therapeutic protein purification"

[γ] Current author affiliation is Center for Precision Genome Editing and Genetic Technologies for Biomedicine, Engelhardt Institute of Molecular Biology, Russian Academy of Sciences, Moscow, Russia.

[¥] Current author affiliation is Radcliffe Department of Medicine, University of Oxford, UK.

### Supporting Information

### S1. Construct schematic and sequence verification

The following is additional detail of the construct schematic and sequence verification performed. Figure S1 displays an illustration of the recombinant *Turnip vein clearing virus* genome prepared as a T-DNA insertion for transformation into *Agrobacterium tumefaciens*. Table S1 displays the primer set used and Figure S2 displays the PCR-based DNA sequence confirmation first in *Escherichia coli* and subsequently in *A. tumefaciens*.

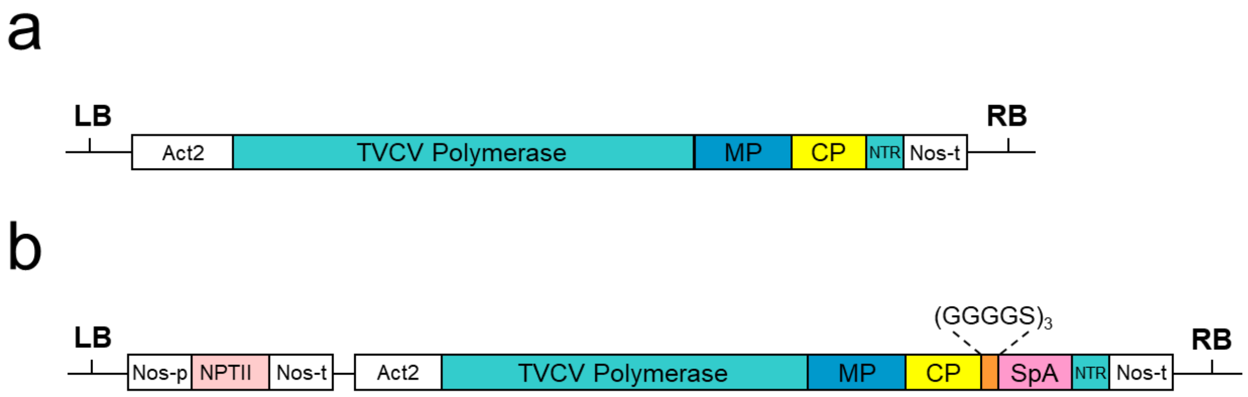

**Figure S1.** *Agrobacterium tumefaciens* T-DNA vector constructs for (a) wild-type TVCV (not used; shown for reference), and (b) TVCV-based immunosorbent nanoparticle. Act2, actin promoter; MP, movement protein; CP, coat protein; NTR, 3' non-translatable region; Nos-p (-t), nopaline synthase gene promoter (terminator); NPTII, kanamycin resistance gene; SpA, Staphylococcal protein A-based Fc-affinity ligand; LB, left border of T-DNA; RB, right border of T-DNA.

**Table S1.** Forward and reverse primers used to verify the vector construct DNA sequence after the bacterial transformation events. The set of primers was designed to generate an amplicon (922 nt) that spans from the start of the coat protein to the end of the Fc affinity ligand.

| Primer | Length (bp) | Sequence |
| --- | --- | --- |
| Forward | 30 | ATGTCTTACAACATTACAAACCCGAATCAG |
| Reverse | 30 | TTCACTCCTTGTTAAAGTTGTTATCTGCCT |

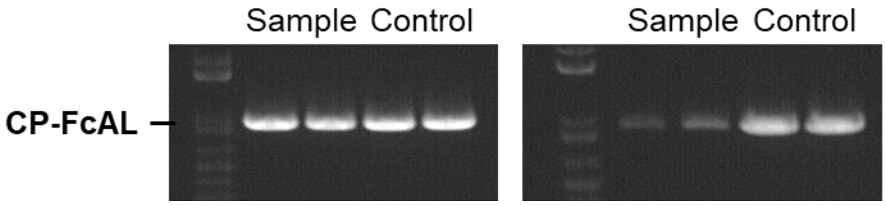

**Figure S2.** DNA gel electrophoresis image of PCR results verify presence of the coat protein Fc-affinity ligand nucleic acids in the transformation of *E. coli* (left) and *A. tumefaciens* (right). The synthesized template DNA is used as control.

#### S2. Mass spectroscopy of coat protein fusion

The following is information on the protein sequence verification performed. Figure S3 shows comparison of the amino acid sequence predicted from the plasmid map (termed control) aligned with the submitted gel electrophoresis band cut VIN sample (termed sample). The sample data was generated by mass spectrometry analysis performed by the UC Davis Proteomics Core (https://proteomics.ucdavis.edu/). Mass spectroscopy data was analyzed using the software Scaffold 4.0 (Proteome Software, Inc., Portland, Oregon).

The mass spectrometry results yielded 78% amino acid coverage (238/306) and 2,203 total spectra at 99.0% minimum with a protein threshold of 5.0% false discovery rate (FDR) and protein decoy FDR of 0.6%, and a peptide threshold of 1.0% FDR and peptide decoy FDR of 0.06%. Results include full coverage of both the N- and C-terminus sequence of the fusion protein (CP + Protein A).

**CP (N-terminal)** **Surface-exposed loop**

Control: MSYNITNPNQYQYFAAVWAEPIPMLNQCMSALSQSYQTQAARDTVRQQFSNLLSAVVTPSQRFPDTGS
Sample: MSYNITNPNQYQYFAAVWAEPIPMLNQCMSALSQSYQTQAAR - - - - QQFSNLLSAVVTPSQRFPDTGS

**TVCV Coat Protein**

Control: RVYVNSAVIKPLYEALMKSFDTRNRIIETEEESRPSASEVANATQRVDDATVAIRSQIQLLLSELSNGHGYMN
Sample: RVYVNSAVIKPLYEALMKSFDTRNRIIETEEESRPSASEVANATQRVDDATVAIRSQIQLLLSELSNGHGYMN

**CP (C-terminal)** **Flexible Linker**

Control: RAEFEALLPWTTAPATGGGGSGGGGSGGGGSGGVTPAANAAQHDEAQQNAFYQVLNMPNLNADQRN
Sample: R - - - - - - - - - - - - - - - - - - - - - - - - - - - - - - - - - - - - - - - - - - - - - - - - - - - - - - - - - - - - - - - - - - N

**Protein A, Domain D + E**

Control: GFIQSLKDDPSQSANVLGEAQKLNDSQAPKADAQQNNFNKDQQSAFYEILNMPNLNEAQRNGFIQSLKD
Sample: GFIQSLKDDPSQSANVLGEAQKLNDSQAPKADAQQNNFNKDQQSAFYEILNMPNLNEAQRNGFIQSLKD

Control: DPSQSTNVLGEAKKLNESQAPKADNNFNKE
Sample: DPSQSTNVLGEAKKLNESQAPKADNNFNKE

**Figure S3.** Mass spectroscopy amino acid analysis coverage of a gel cut VIN sample compared to the plasmid map sequence. CP, coat protein; TVCV, turnip vein clearing virus.

#### S3. Supporting immunosorbence characterization

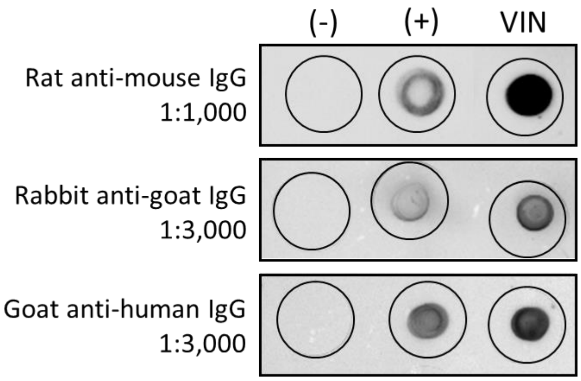

**Figure S4.** Dot blot assay portrays the negative control (-), wild-type TMV, positive control (+), recombinant Protein A, and VIN binding to Immunoglobulin G from rat, rabbit, and goat. The assay only used secondary antibodies, given the nature of the immunosorbent probe.

The following is functionality verification of VINs produced via mechanical transmission. Figure S5 depicts the bind-and-elute operation performed with VIN produced via mechanical inoculation of *Nicotiana benthamiana* plants. There is a visible difference in the purified VIN solution produced via mechanical transmission, in what we suspect is a discernable increase in coat protein degradation products given by the band at ~25 kD.

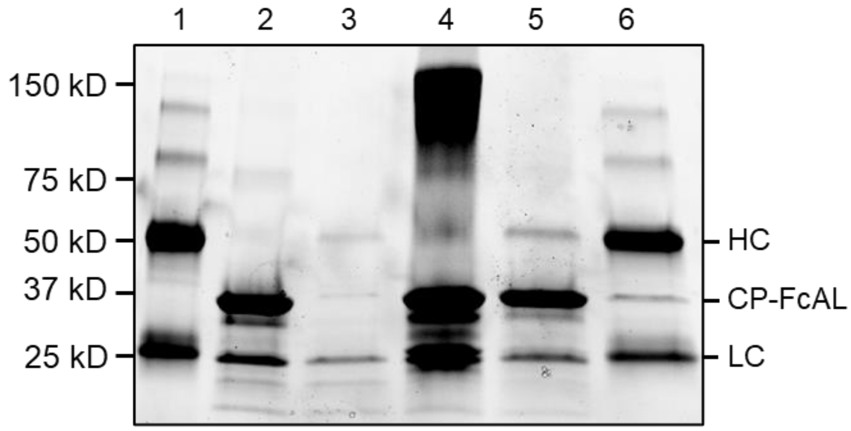

**Figure S5.** SDS-PAGE of mechanical transmission-generated VIN bind-and-elute with human immunoglobulin G. Description of lanes are 1 – initial human antibody, 2 – initial VIN, 3 – centrifuge supernatant (waste stream), 4 – resuspended pellet, 5 – recovered VIN, 6 – recovered human antibody. CP, virus coat protein; FcAL, Fc-affinity ligand; HC, antibody heavy chain; LC, antibody light chain.

We observed that centrifugations of 20,000 x g for 90 minutes at 4 °C were sufficient to pellet the VINs in the absence of a suitable population of target protein to bind with, as demonstrated in Figure S6 by the use of bovine serum albumin (BSA) as a negative control target protein substitution and the complete diversion of BSA to the supernatant loss lane.

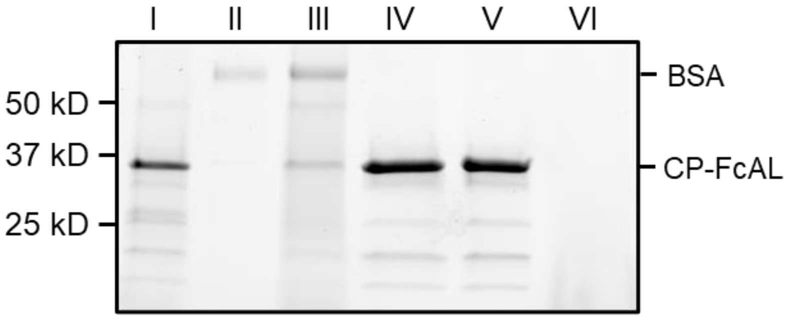

**Figure S6.** SDS-PAGE results of the VIN-based bind-and-elute procedure with a target of bovine serum albumin (BSA) using a centrifugation condition of 20,000 x g for 90 minutes at 4 °C. Lane definitions: I – initial VIN added; II – initial BSA added; III – VIN/BSA supernatant (loss); IV – VIN/BSA pellet resuspended; V – recovered VIN; VI – recovered BSA from resuspended pellet.

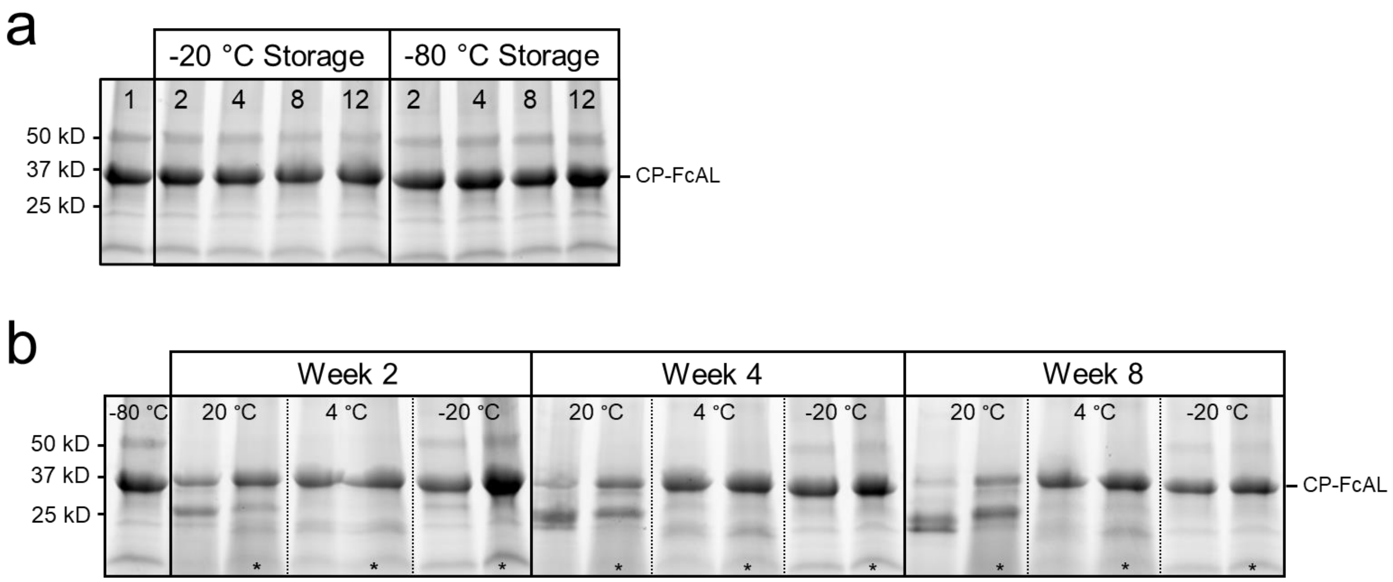

**Figure S7.** (A) The effect of freeze-thaw (FT) cycles (2, 4, 8, 12) on the stability of the VIN coat protein Fc-affinity ligand fusion using both -20 °C and -80 °C storage conditions shown by SDS-PAGE. The initial stock (FT cycle 1) is preserved in -80 °C storage. (B) Long-term storage stability of the VIN coat protein Fc-affinity ligand fusion at 20 °C, 4 °C, and -20 °C temperature conditions with and without protease inhibitor additives. Lanes with an asterisk (*) have 2 mM EDTA and 1 mM PMSF protease inhibitors added to the storage solution.

#### S4. Process development study results

The following is additional detail that highlights some of the key experiments performed as part of the VIN purification and use process development. Figure S7 shows the results the of 2-factor 2-level study design for extraction optimization of the VIN purification. Figure S8 displays the results of a temperature screen performed for various VIN and wild-type *Tobacco mosaic virus* (wt-TMV) solutions as part of a thermostability assessment. Figure 9 displays the bind-and-elute assessment which was used to confirm retention of functionality for the different VIN preparations.

Interestingly, the addition of protease inhibitors to the extraction buffer resulted in increased losses to VIN recovery. Corresponding with other published studies, the use of a low pH extraction buffer improved impurity clearance and thus VIN purity. The temperature screen shows the difference in apparent stability and/or presence of VIN in solution compared between the final VIN solution and the crude post-extraction solution, illuminating an unexpected instability of the VIN in crude solution. This difference in temperature screen results between the crude and purified solution was not observed for wt-TMV. The authors did not extend this study further to assess whether this can be attributed to impurity-enhanced degradation or sedimentation of the VIN.

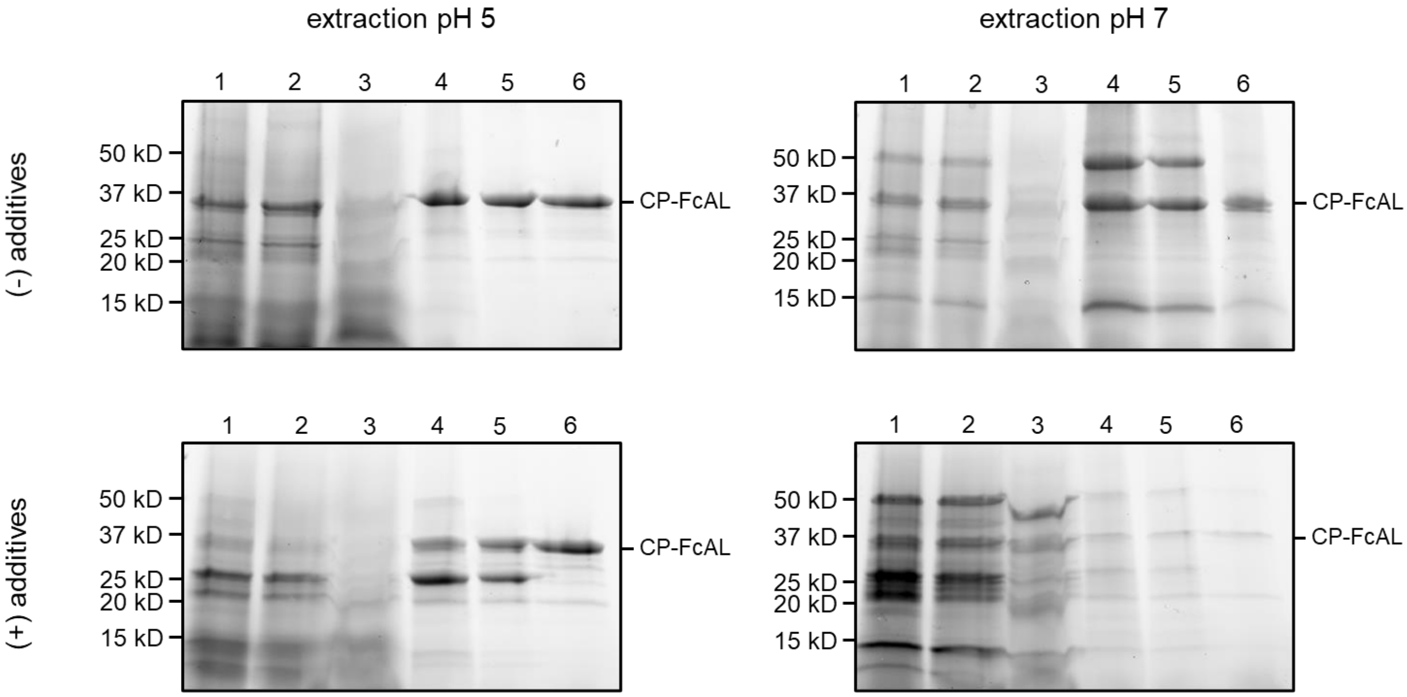

**Figure S8.** Process development observations via SDS-PAGE for a 2-factor 2-level study design to test extraction conditions (buffer pH and buffer additives). Buffer additives are 2 mM EDTA and 1 mM PMSF. Lane definitions: 1 – crude extraction; 2 – clarification; 3 – PEG precipitation (supernatant); 4 – PEG precipitation (pellet); 5 – post-PEG centrifugation (supernatant); 6 – final VIN solution.

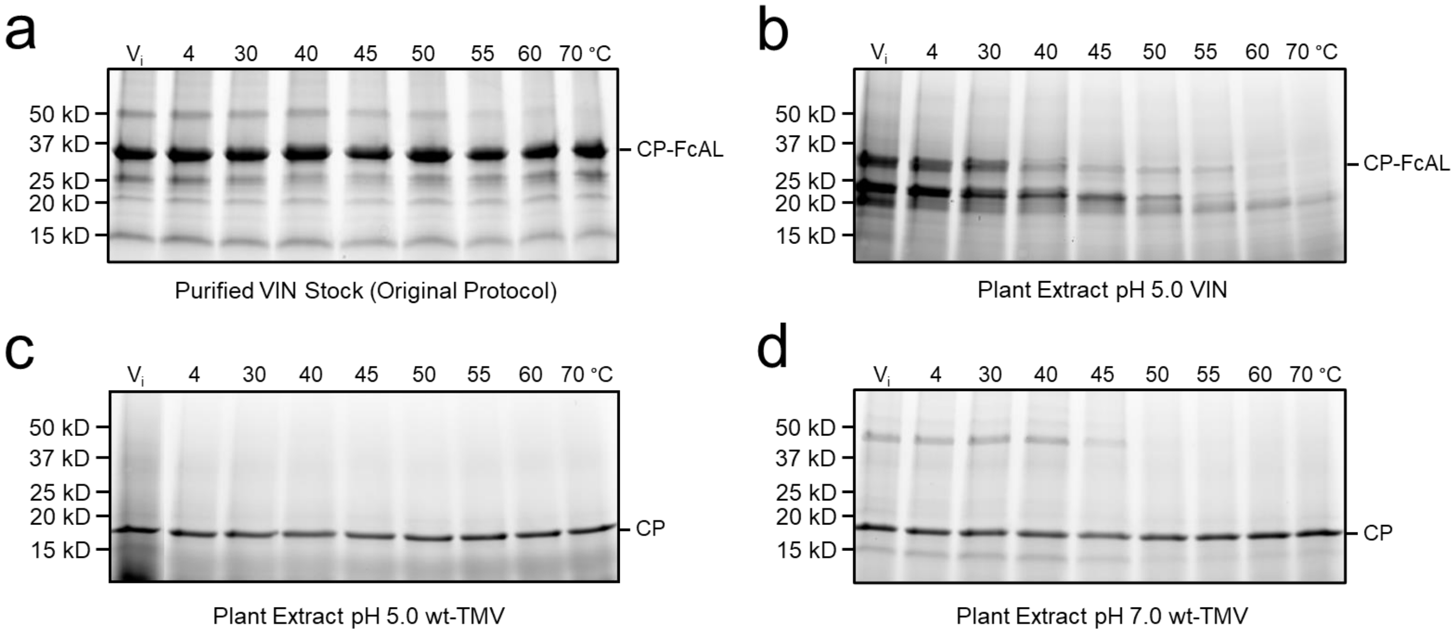

**Figure S9.** Key thermostability assessment results visualized with SDS-PAGE after five minute temperature holds ranging from 4 °C to 70 °C of (A) VIN purified using the original protocol, (B) VIN in crude plant extract at pH 5.0 with additives, (C) wt-TMV in crude plant extract at pH 5.0 with additives, and (D) wt-TMV in crude plant extract at pH 7.0 with additives. Samples are lightly centrifuged (5,000 x g for 5 minutes at 4 °C) after heat hold to clear aggregates from suspension. $V_i$, initial virus solution.

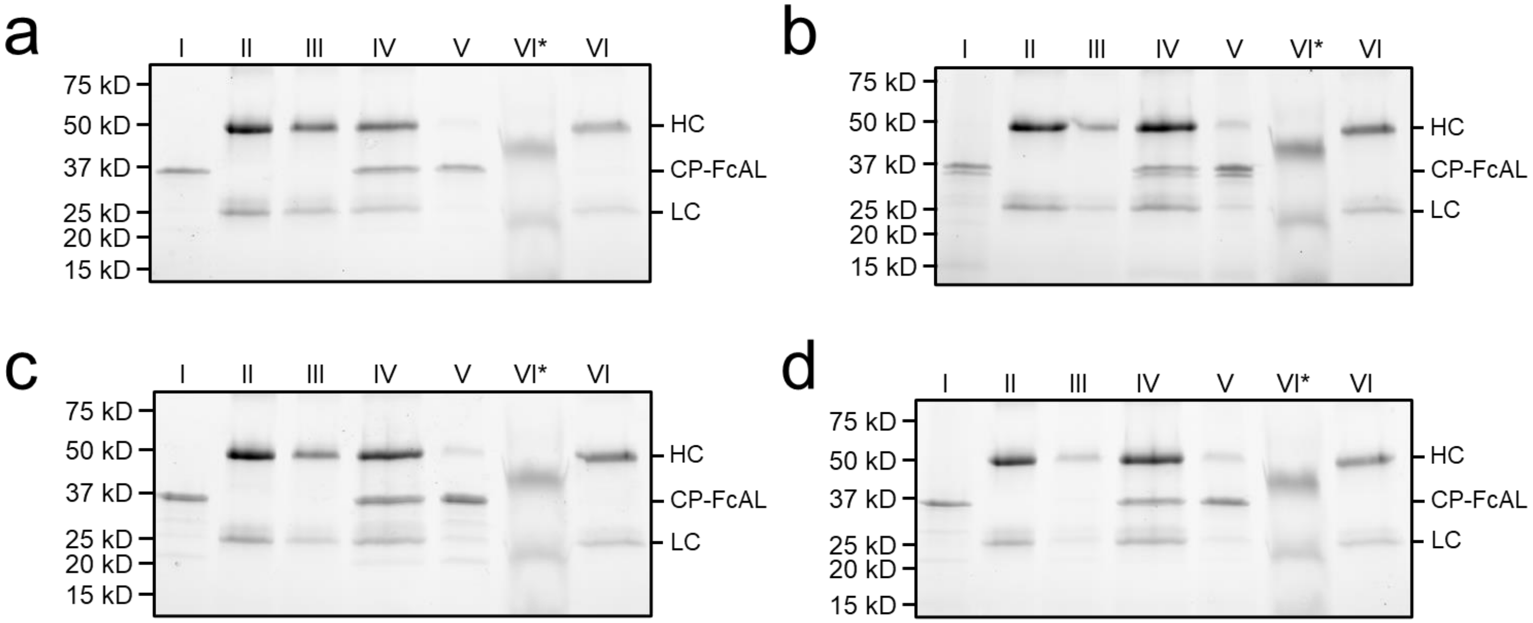

**Figure S10.** Performance verification of VIN bind-and-elute functionality for those generated through the three most promising process improvement workflows of (a) low pH extract with a heat hold, (b) neutral pH extraction with a heat hold, (c) low pH extract plus buffer additives with heat hold, and (d) the original protocol methodology. Lanes are labeled according to the naming convention outlined in Figure 4a with the addition of VI*, which is the intermediary step containing PEG 6,000 prior to target protein buffer exchange.
